# Investigation of the impact of stool collection methods on the metabolomics analysis/profiles of infant fecal samples

**DOI:** 10.1101/2021.09.21.461251

**Authors:** Chiara-Maria Homann, Sara Dizzell, Sandi M. Azab, Eileen K. Hutton, on behalf of the GI-MDH Consortium, Katherine M. Morrison, Jennifer C. Stearns

## Abstract

Metabolomic studies are important to understand microbial metabolism and interaction between the host and the gut microbiome. Although there have been efforts to standardize sample processing in metabolomic studies, infant samples are mostly disregarded. In birth cohort studies, the use of diaper liners is prevalent and its impact on fecal metabolic profile remains untested. In this study, we compared metabolite profiles of fecal samples collected as solid stool and those collected from stool saturated liner. One infant’s stool sample was collected in triplicate for solid stool and stool saturated liner. Comprehensive metabolomics analysis of the fecal samples was performed using NMR, UPLC and DI-MS. The total number, identities and concentrations of the metabolites were determined and compared between stool sample collection methods (stool vs. liner). The number and identity of metabolites did not differ between collection methods for NMR and DI-MS when excluding metabolites with a coefficient of variation (CV) > 40%. NMR analysis demonstrated lowest bias between collection methods, and lowest technical precision between triplicates of the same method followed by DI-MS then UPLC. Concentrations of many metabolites from stool and stool saturated liner differed significantly as revealed by Bland-Altman plots and t-tests. Overall, a mean bias of 10.2% in the Bland-Altman analysis was acceptable for some metabolites confirming mutual agreement but not for others with a wide range of bias (−97-117%). Consequently, stool and stool-saturated liner could be used interchangeably only for some select metabolite classes e.g. amino acids. Differences between the metabolomic profiles of solid stool samples and stool saturated liner samples for some important molecules e.g., ethanol, fumarate, short chain fatty acids and bile acids, indicate the need for standardization in stool collection method for metabolomic studies performed in infants.

## 1. Introduction

The microorganisms that reside within the human gut intimately interact with the host - immunologically, and metabolically [1]. Methods to study the gut microbiome include DNA and RNA based methods that provide information about microbial genes and pathways; however, these methods can only predict microbial and host metabolism. In order to study metabolic differences associated with health and disease, microbial and host metabolites can be measured directly using metabolomics. Metabolomics is the high-throughput identification and quantification of small molecules in body tissue or biofluids [2]. It is one of the newer -omics technologies/disciplines [3]. Gut metabolites are very diverse and many of them remain uncharacterized. As such, a number of different methods have emerged to resolve and quantify the constituents and diversity of the gut metabolome.

The gut microbiota is often explored in relation to the fecal metabolite profile and there is high interrelation between the gut microbiome and the fecal metabolome [4]. The fecal metabolome has been said to provide a “complementary functional readout” of microbial metabolism as well as the interaction between the gut microbiome and the host [4]. The relationship between metabolites and the microbiota within the human gut has been explored in adults [5-7]; however, less is known about the gut metabolome in infants. Establishment of the infant gut microbiome occurs in the first few years of life, which is a critical time in development and is emerging as an important predictor of later health outcomes [8-10]. Currently, the most common analysis of the gut microbiome in infants is performed using culture independent methods, such as shotgun metagenomic sequencing or taxonomic profiling with use of the 16S rRNA marker gene. Standard protocols are important for reproducibility in research and within the microbiome field some important work has been done to document the variation that arises due to technical and methodological differences between studies and study centers [11-14].

Standard protocols exist to collect adult [15] and infant stool for nucleic acid analysis, but whether these same collection methods are appropriate for metabolites is not known. There is a lack of standardization in metabolomics study protocols and sample collection methods, especially for fecal metabolomics analysis [3]. Additionally, studies that investigate methodological differences and their implications have solely been performed using adult samples [3]. Infants produce stool of varying consistencies and breastfeeding babies in particular often do not have solid stool, making sample collection challenging. One of the approaches to the challenge of infant stool samples has been implementing the use of a standard diaper liner to collect stool residue if solid stool is not available. Therefore, it is important to determine if similar metabolite profiles, in terms of the type and number of unique metabolites, are obtained when different methods of sample collection are used, namely from solid stool or stool saturated diaper liner. In this paper we compared these two collection methods from one stool sample in triplicate across three metabolomic platforms (nuclear magnetic resonance, NMR; direct infusion mass spectrometry, DI-MS, and ultra-high performance liquid chromatography, UPLC) to determine the metabolic and technical variation introduced by the collection method.

## 2. Results

### 2.1. Metabolite Coverage

After removing metabolites with coefficient of variation (CV) > 40%, 159 unique metabolites were detected altogether in the solid stool and stool saturated liner samples (Figure 1A). 65 metabolites were detected by NMR, 79 by DI-MS and 39 by UPLC. While none of the metabolites detected by UPLC were measured with the other platforms, 24 metabolites were measured by both NMR and DI-MS (Figure 1A). The overlapping metabolites included mostly amino acids e.g., alanine, asparagine, isoleucine, leucine, and valine., saccharides e.g., glucose, and short-chain fatty acids e.g., acetate, butyrate, and propionate. To assess the variation of metabolites within a sample for the two sample collection methods, the chemodiversity index was calculated for each sample based on the number of unique metabolites across all three metabolomic methods. The chemodiversity index did not differ significantly between sample collection methods (p = 0.32), which indicates that the same metabolites are found in solid stool samples and stool saturated liners. This is further confirmed by the number of metabolites found with each sample collection method and metabolic platform, where metabolite overlap between stool collection methods ranges from 95.0 - 100% (Table 1, Figure 1A), prior to and after the exclusion of metabolites with CV > 40%. A Bland-Altman plot was generated to assess overall mutual agreement between sample collection methods (Figure 1B). Although, the mean bias between stool collection methods was 10.2% (p = 0.019) for 159 metabolites, the 95% confidence interval was wide, indicating that concentrations differed by stool collection method and suggesting that stool saturated liner samples had average metabolite concentrations that were lower than average metabolite concentrations in solid stool samples. Additionally, a few metabolites were outside of the limits of agreement for all metabolomic platforms (indicated by arrows in Figure 1B), signifying that there were drastic differences in the concentrations for some metabolites e.g., ethanol, formate, and fumarate, creatinine and arginine based on stool sample collection method.

**Figure 1.**
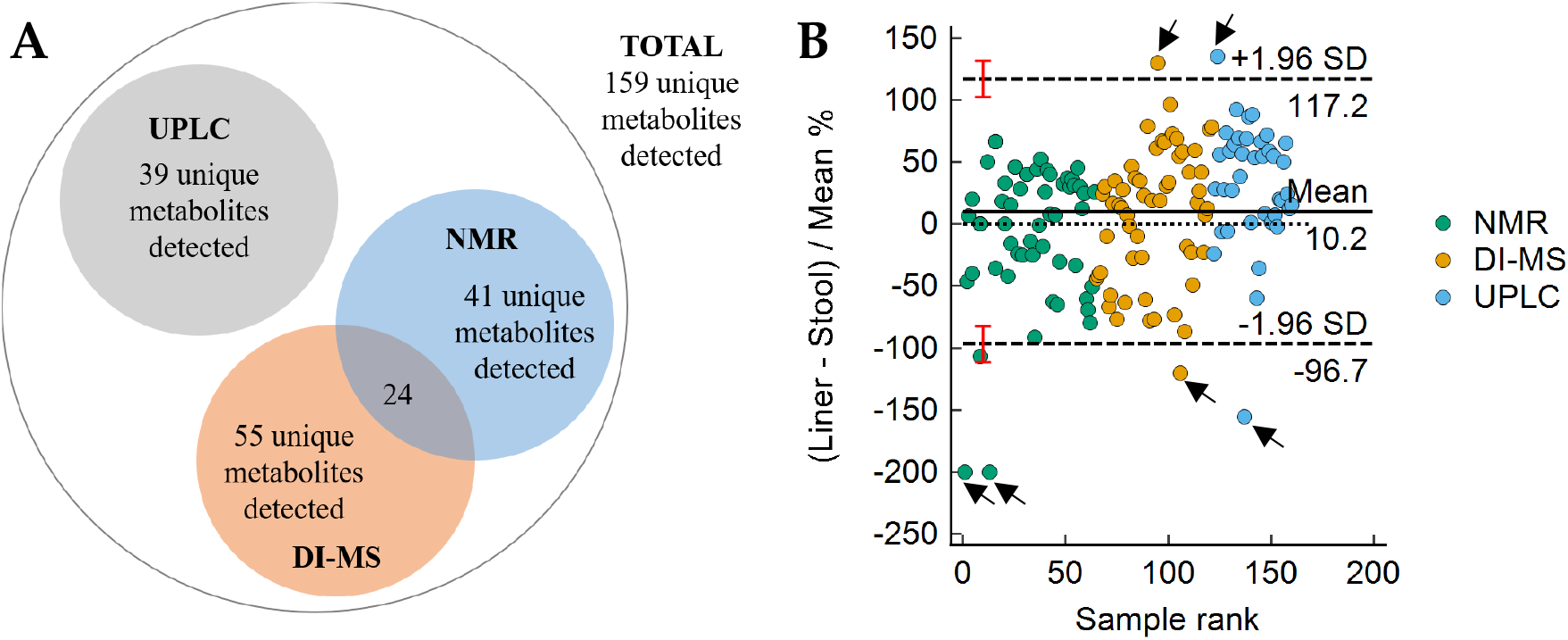
(**A**) Number of metabolites measured per platform (UPLC, NMR, DI-MS) after metabolites with a coefficient of variance greater than 40% were excluded; (**B**) Bland-Altman plot comparing metabolite concentrations between methods for stool collection (solid stool vs. liner), based on sample rank and colored by metabolic platform. Metabolites outside of the confidence intervals of the upper and lower limits are considered different between methods and are indicated by the black arrows.

**Table 1.** Comparison of the number of metabolites measured using different metabolomic methods and collection methods.

|  | Number of metabolites measured (n) | Number of metabolites detected (n) | Number of metabolites detected in solid stool (n, %) | Number of metabolites detected in liner (n, %) | Overlap between solid stool and liner (n, %) |
| --- | --- | --- | --- | --- | --- |
| **No CV cut-off** |  |  |  |  |  |
| NMR | 67 | 67 | 67 (100) | 67 (100) | 67 (100) |
| DI-MS | 116 | 110 | 109 (99.1) | 108 (98.2) | 107 (97.3) |
| UPLC | 68 | 40 | 40 (100) | 38 (95.0) | 38 (95.0) |
| **CV < 40%** |  |  |  |  |  |
| NMR | 67 | 65 | 65 (100) | 65 (100) | 65 (100) |
| DI-MS | 116 | 79 | 79 (100) | 79 (100) | 79 (100) |
| UPLC | 68 | 39 | 39 (100) | 37 (95.0) | 37 (95.0) |

### 2.2. Metabolites Measured with Nuclear Magnetic Resonance (NMR)

Quantitative NMR spectroscopy was used for targeted metabolomic analysis of water-soluble metabolite classes including amino acids, saccharides, alcohols, organic acids, amines, tricarboxylic acid (TCA) cycle intermediates, and short chain fatty acids (SCFAs). A total of 67 metabolites were measured and 65 were detected in both solid stool and stool saturated liner samples (CV < 40%) (Table 1). Ethanol, formate, and fumarate were outside of the limits of agreement of the Bland-Altman plot (Figure 1B), indicating significant differences between sample collection methods. Ethanol and fumarate had higher concentrations in solid stool samples, whereas creatinine had higher concentrations in stool saturated liner samples. In further analyses with paired t-tests, the concentrations of 56 metabolites were significantly different between stool saturated liner and solid stool samples after adjustment for multiple testing (Figure S2A). Of the physicochemical characteristics, only polar surface area was significantly associated with the percentage difference in concentrations between the stool sample collection methods. Higher polar surface area was associated with higher metabolite concentrations in stool saturated liner samples (β = −0.17, p = 0.017). Metabolite characteristics such as polarizability, solubility in water, molecular weight, or physiological charge were not associated with the difference in concentration between solid stool and stool saturated liner (Figure S2B, p > 0.05). Variation between technical replicates was lower for metabolite measurements by NMR compared with the variation seen for the other metabolic platforms. Mean (SD) variation for solid stool samples was 2.8% (SD = 2.50), and average variation for stool saturated liner samples was 3.4% (SD = 2.87) (Table 2).

**Table 2.** Coefficients of variation by metabolic platform after metabolites with coefficients of variations > 40% were excluded.

|  | Solid Stool |  |  | Diaper Liner |  |  |
| --- | --- | --- | --- | --- | --- | --- |
|  | Mean (SD) | Median | Range (Min, Max) | Mean (SD) | Median | Range (Min, Max) |
| NMR | 2.81 (2.50) | 2.22 | 12.32 (0.059, 12.38) | 3.37 (2.87) | 2.57 | 16.19 (0.19, 16.39) |
| DI-MS | 10.03 (9.62) | 8.13 | 34.54 (0.00, 34.60) | 8.75 (8.79) | 7.51 | 34.64 (0.0, 34.64) |
| UPLC | 15.48 (6.66) | 14.43 | 31.36 (6.19, 37.55) | 17.06 (5.52) | 16.58 | 26.85 (5.88, 32.73) |

### 2.3. Metabolites Measured with Direct Flow Injection Mass Spectrometry (DI-MS)

For targeted metabolomic analysis of biogenic amines, amino acids, acylcarnitines, phospholipids and sphingolipids, direct flow injection mass spectrometry (DI-MS) was used. A total of 116 metabolites were measured and 79 of these metabolites were detected in both solid stool and stool saturated liner samples after the CV cut-off (Table 1). For this metabolic platform, only one of the metabolites, arginine, was outside of the limits of agreement in the Bland-Altman plot (Figure 1B). Significant differences were found in the concentration of DI-MS measured metabolites in stool saturated liner and solid stool samples in paired t-tests after adjustment for multiple testing for 16 metabolites (Figure S3A). There was noticeable technical variation with this method, however, which reduced precision of these measurements. Mean (SD) CV for solid stool samples was 10.0% (SD = 9.62) and 8.8% (SD = 8.79) for stool saturated liner samples (Table 2). None of the metabolite characteristics were associated with the difference in metabolite concentrations between solid stool and stool saturated liner samples (Figure S3B, p > 0.05).

### 2.4. Metabolites Measured with Ultra-High Performance Liquid Chromatography (UPLC)

UPLC was used as a targeted metabolomic method to examine bile acids in the stool samples. A total of 68 metabolites were measured, however, only 39 metabolites were retained after the CV cut-off; all 39 were detected in solid stool and 37 metabolites were detected in stool saturated liner (Table 1), Glycochenodeoxycholic acid and isolithocholic acid were not detected in stool saturated liner. No metabolite had a significantly higher or lower concentration within solid stool versus stool saturated liner samples in paired t-tests after adjustment for multiple testing (Figure S4A). As expected, the concentration of the metabolites measured with UPLC was ten-fold lower than with the other methods, with many metabolites recorded near the limit of detection due to their low abundance, and there was more technical variation than with NMR which may have reduced precision. Mean (SD) of CV for solid stool samples was 15.5% (6.66) and 17.1% (5.52) for stool saturated liner samples (Table 2), which were the highest average CVs of the three metabolic platforms. As previously observed for DI-MS, metabolite characteristics were not associated with their concentration measured from solid stool or stool saturated liner samples (Figure S4B, p > 0.05).

### 2.5. Short-Chain Fatty Acids

One of the metabolite groups of higher interest were the SCFAs, as these are known intermediates and end-products of bacterial metabolism. Paired t-tests were performed to analyze the differences in absolute concentrations between sample collection methods. The concentrations of all the SCFAs, as well as their total concentration differed significantly by sample collection method. Acetate (p = 0.011), butyrate (p = 0.00084), propionate (p = 0.0016), isovalerate (p = 0.0027), valerate (p = 0.00025) and total SCFAs (p = 0.0066) had higher concentrations in the stool saturated liner samples, while isobutyrate (p = 0.0011) had a higher concentration in the solid stool sample (Table 3).

**Table 3.**
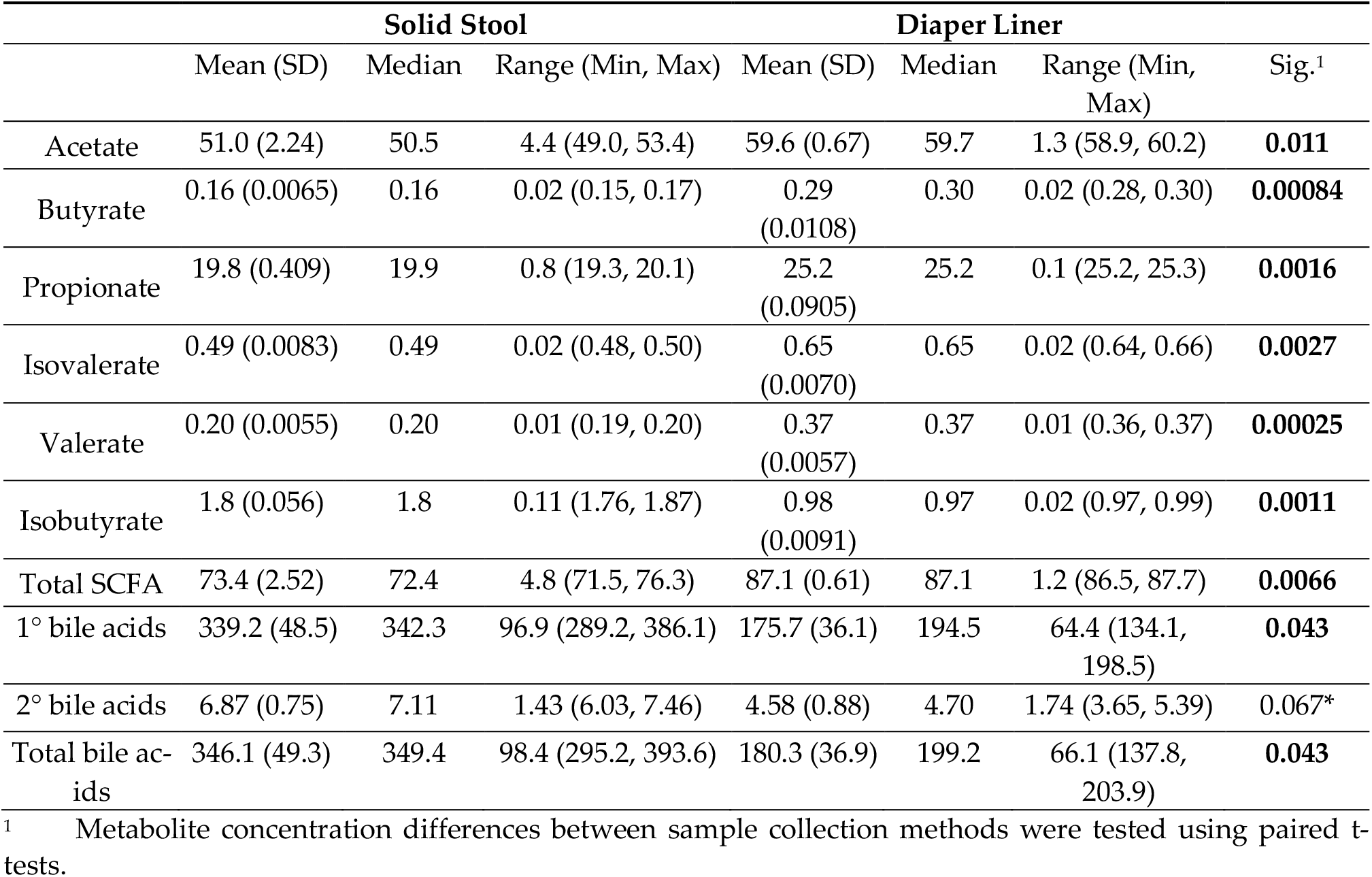
Concentrations of the metabolites of interest: SCFAs (µmol/g) and bile acids (nmol/g).

### 2.6. Bile Acids

Another metabolite group of interest were the 15 major bile acids, as microbes are responsible for the conversion of primary bile acids to secondary bile acids. These are cholic acid, chenodeoxycholic acid, taurocholic acid, taurochenodeoxycholic acid, glycocholic acid, glycochenodeoxycholic acid, lithocholic acid, deoxycholic acid, ursodeoxycholic acid, glycolithocholic acid, taurolithocholic acid, glycodeoxycholic acid, taurodeoxycholic acid, glycoursodeoxycholic acid and tauroursodeoxycholic acid. Total concentrations for the primary, secondary, and total bile acids were calculated and compared between sample collection methods with paired t-tests. Primary bile acid concentrations and total bile acid concentrations were significantly higher in solid stool samples than in stool saturated liner samples (p = 0.043; p = 0.043). Secondary bile acids were trending towards a significantly higher concentration in solid stool samples (p = 0.067) (Table 3).

## 3. Discussion

Collection and banking of stool samples from current or previous cohorts for later metabolomic analysis is an important activity. It is important to strike a balance between ease of sample collection and the comparability of results across collection methods, especially in breastfeeding infants, where stool consistency can provide a challenge for sample collection. In our study, Baby, Food & Mi, we have several infants for which only stool saturated diaper liner was available. Undertaking an extensive study of fecal metabolites measured using three different analytical techniques (NMR, DI-MS and UPLC) described here, we sought to explore whether metabolomic profiles of solid stool samples are comparable to those found with stool saturated liner. In this comparative study, we showed that after a cut-off for high technical variation, the individual metabolites detected in solid stool samples and stool saturated liner did not differ between sample collection methods; however, metabolite concentrations differed significantly between collection methods when analyzed on all three metabolomic platforms: NMR, DI-MS and UPLC. Discrepancies were also seen for metabolite groups of specific interest, namely SCFAs and bile acids. These metabolite groups in particular showed differences in concentrations between solid stool and stool saturated liners, where all SCFAs except isobutyrate, had higher concentrations in stool saturated liner samples and bile acids had higher concentrations in solid stool samples, indicating that the sample collection method is an important consideration in infant metabolomic studies.

Metabolomic analysis performed in this study was based on two analytical systems: nuclear magnetic resonance and mass spectrometry. NMR is a well-established platform [16] and results are highly reproducible, inherently quantitative, robust, and cost effective [3,17,18]. However, NMR is less sensitive than mass spectrometry, by a factor of 10 - 100x, and therefore, has a narrower coverage of metabolites [3,18]. Reduced technical variation between samples from each collection method could surely impact the statistical power of the t-tests performed and the resulting findings across all analytical platforms with an advantage towards NMR that exhibits highest precision. As a high-throughput method, the advent of mass spectrometry introduced a new dimension to medical research. Two mass spectrometry methods were described in this study: UPLC-MS/MS and DI-MS. UPLC is a separation-based method, while DI-MS is separation free [19]. MS methods are more sensitive than NMR and are often targeted for specific metabolites [17]. However, MS methods have less reliable molecule quantification, need internal standards and are more prone to error due to matrix effects [3,17]. Though not performed in this study, there is a high variety of MS-based methods for metabolomic analysis including GC-MS, CE-MS and MALDI-MS; many studies primarily use NMR and GC-MS [19].

Benchmarking the effect of the collection method (solid stool vs. stool saturated liner) on the resulting metabolite concentrations is essential prior to undertaking large-scale analyses. This is especially important in large birth cohort studies with breastfeeding infants, where the use of diaper liners is prevalent, and stool often does not have a solid consistency. Factors affecting the metabolite profile in fecal samples are sample collection methods, sample storage, as well as sample preparation [3]. Previous studies have indicated that freezing does not affect the metabolic profile of stool samples. The suggested workflow for fecal metabolomic samples is to keep fresh samples on ice until they reach the laboratory and can be processed. After initial processing samples should be kept at −80 degrees Celsius before chemical analysis starts [2]. Multiple freeze-thaw cycles should be avoided since it could alter stool metabolite profiles [20]. The benefit of freezing samples is that preservatives are not needed [15]. Multiple studies have investigated the use of preservatives in fecal samples, assessing stability of the stool sample at room temperature. These studies have shown that preservation in 95% ethanol shows the highest concordance with samples frozen quickly after collection - also considered the gold standard for fecal sample collection [15,21]. In our study, however, stool sample preservatives were not used, as sample was frozen quickly after defecation, following the evidence-backed protocol outlined above. Other factors influencing fecal samples include stool water content, which was not considered here.

Sample collection is an important consideration in metabolomics. To our knowledge, this study is the first of its kind investigating differences between method of stool sample collection (solid stool vs. stool saturated liner) in infant samples, and its impact on the metabolomics profiles. Therefore, this study provides valuable insight into differences in technique for analyzing the metabolites of infant fecal samples. Limitations of this study include the limited sample size (n = 1) and the high technical variation for some metabolites within the three technical replicates for each method, especially for DI-MS and UPLC. Both of these factors reduce the statistical power of the study, which might conceal actual associations when testing biological hypotheses. Thus, to fully understand the methodological differences between solid stool and diaper liner samples this study could be expanded to include more infants. After observing differences in metabolite concentrations for all three of the metabolomic methods, NMR, DI-MS and UPLC, the practical implications of this study are to solely use one of the sample collection methods for metabolomic analysis, preferably solid stool samples, as stool saturated liner failed to detect one of the major bile acids glycochenodeoxycholic acid. Within a targeted approach for studies only interested in the analysis of amino acids, solid stool and stool saturated liner could be used interchangeably.

## 4. Materials and Methods

### 4.1. Participant Recruitment

One infant from the Baby, Food & Mi study [22], a sub-cohort of 15 infants from the primary Baby & Mi study at McMaster University in Hamilton, Ontario, Canada was observed [23].

### 4.2. Fecal Sample Collection

The study participant was asked to use a diaper liner (Bummis, Quebec, Canada) during the sample collection period; samples were collected at around 6 months of age. Fecal samples were collected by placing the infant’s soiled diaper, including the diaper liner in a resealable bag with an anaerobic sachet (Fisher Scientific, Hampshire, UK) immediately after the infant defecated. The bag containing the sample was then placed in an insulated cooler bag with a frozen reusable ice pack and transported to McMaster University. Processing of the sample occurred in a Bactron IV anaerobic chamber (Sheldon Manufacturing INC, Cornelius, OR). Solid stool aliquots of 100 mg, 50 mg, 100 mg, and 600 mg were measured and aliquoted into cryovials for DNA isolation, Direct Flow Injection Mass Spectrometry (DI-MS), Quantitative Nuclear Magnetic Resonance Spectroscopy (NMR) and ultra-performance liquid chromatography-tandem mass spectrometry (UPLC), respectively. Stool saturated liner samples containing 100 mg, 50 mg, 100 mg, and 600 mg of stool were cut from the liner and also aliquoted into cryovials. To ensure that the amount of solid stool in the stool saturated liner samples was accurate, a clean liner sample with no stool was weighed and subtracted from the weight of the stool saturated liner. Clean liner was used as a negative control for metabolite extraction. All samples (solid stool, stool saturated liner and clean liner) were aliquoted in triplicate for each metabolomic method and were stored at −80 °C until further processing could occur. Samples were shipped on dry ice to The Metabolomics Innovation Centre (TMIC; Alberta, Canada) where metabolic profiling was done according to standard protocols, described briefly below.

### 4.3. Metabolomic Profiling

Samples were prepared according to [24] and [25]. Total metabolites were measured with nuclear resonance spectrometry (NMR). All ^1^H-NMR spectra were collected on a 700 MHz Avance III (Bruker) spectrometer equipped with a 5 mm HCN Z-gradient pulsed-field gradient (PFG) cryoprobe. ^1^H-NMR spectra were acquired at 25°C using the first transient of the NOESY presaturation pulse sequence (noesy1dpr), chosen for its high degree of quantitative accuracy. All free induction decays were zero-filled to 250 K data points. The singlet produced by the DSS methyl groups was used as an internal standard for chemical shift referencing (set to 0 ppm). All ^1^H-NMR spectra were processed and analyzed using the Chenomx NMR Suite Professional software package version 8.1 (Chenomx Inc., Edmonton, AB). Untargeted metabolomics was performed with direct flow injection mass spectrometry with an Agilent 1100 series HPLC system (Agilent, Palo Alto, CA) and an Agilent reversed-phase Zorbax Eclipse XDB C18 column (3.0 mm × 100 mm, 3.5 µm particle size, 80 Å pore size) with an AB SCIEX QTRAP® 4000 mass spectrometer (AB SCIEX, CA, U.S.A.). The controlling software was Analyst® 1.6.2. The mass spectrometer was set to positive electrospray ionization with multiple reaction monitoring (MRM) mode. Bile acids were measured with Ultra-Performance Liquid Chromatography-Tandem Mass Spectrometry (UPLC) on an Agilent 1290 system coupled to a 4000 QTRAP mass spectrometer. The MS instrument was operated in the multiple-reaction monitoring (MRM) mode with negative-ion (-) detection. A Waters BEH 15-cm long, 2.1-mm I.D. and C18 LC column was used, and the mobile phase was (A) 0.01% formic acid in water and (B) 0.01% formic acid in acetonitrile for binary-solvent gradient elution by RPLC. Linear regression calibration curves were constructed between analyte-to-internal standard peak area ratios (As/Ai) versus molar concentrations (nmol/mL).

### 4.4. Statistical Analysis

Data analysis was performed in R [26] and MedCalc [27]. Metabolites with coefficient of variation (CV) greater than 40% were excluded from statistical analysis. MedCalc software was used to generate a Bland-Altman plot, comparing concentrations of the metabolites between solid stool and stool saturated liner for the metabolomic platforms. For metabolites that were measured with two metabolomic platforms, the metabolomic platform with the lower CV for the metabolite was included in the Bland-Altman plot. Statistical significance between solid stool and stool saturated liner samples was determined with two-tailed paired t-tests with FDR-adjustment for multiple testing. Physical and chemical properties for each metabolite were obtained from the Human Metabolome Database [28] and included solubility in water, physiological charge, polarizability, polar surface area and molecular weight. Associations of metabolite characteristics with the percentage difference in concentration between solid stool and stool saturated liner were tested using univariate linear regressions with the stats package [26]. The chemodiversity index is an alpha diversity measure that describes the variation of metabolites within a sample [29,30] and was calculated for each sample:

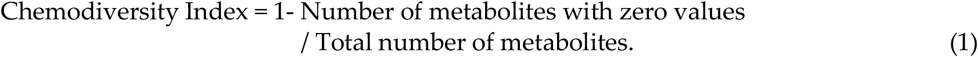

Differences in chemodiversity index by sample collection method were calculated with a two-tailed paired t-test. For the investigation of specific metabolite groups (SCFAs and bile acids), individual and total SCFA concentrations, as well as primary, secondary, and total bile acid concentrations were calculated and compared between sample collection methods using two-tailed paired t-tests. The cut-off point for all statistical analyses presented here is p < 0.05.

## 5. Conclusions

This study establishes that there is an association between stool sample collection method and metabolite profiles in three common metabolomic analysis methods, namely nuclear magnetic resonance, direct infusion mass spectrometry and ultra-high performance liquid chromatography. This highlights the need to either standardize research protocols to one of the stool collection methods or to control for stool collection method in analyses.

## Supplementary Materials

The following are available online at www.mdpi.com/xxx/s1, Table S1: CV for metabolites measured in stool and liner with NMR, Table S2: CV for metabolites measured in stool and liner with DI-MS/MS, Table S3: CV for metabolites measured in stool and liner with UPLC-MS, Figure S1: Bland-Altman plot comparing metabolite concentrations between methods for stool collection (solid stool vs. liner), colored by metabolic platform. Metabolites outside of the confidence intervals of the upper and lower limits are considered different between methods, Figure S2: Metabolite, including short chain fatty acid, concentrations (µmol/g) measured in stool and liner with NMR. (**A)** Boxplot of metabolite concentrations, * = significant differences between stool sample collection methods, after adjustment for multiple testing. (**B)** Characteristics of the metabolites in relation to their log2 fold change by sample collection method., Figure S3: Metabolite concentrations (µmol/g) measured in stool and from liner with DI-MS/MS. (**A)** Boxplot of metabolite concentrations, * = significant differences between stool sample collection methods, after adjustment for multiple testing. **(B)** Characteristics of the metabolites in relation to their log2 fold change by sample collection method., Figure S4: Bile acid concentration (nmol/g) measured in stool and from liner with UPLC-MS. (**A)** Boxplot of metabolite concentrations. (**B)** Characteristics of the metabolites in relation to their log2 fold change by sample collection method.

## Author Contributions

Conceptualization, Eileen Hutton, Katherine Morrison and Jennifer Stearns; Data curation, Sara Dizzell; Formal analysis, Chiara-Maria Homann and Jennifer Stearns; Funding acquisition, Eileen Hutton, Katherine Morrison and Jennifer Stearns; Methodology, Sara Dizzell, Sandi Azab and Jennifer Stearns; Project administration, Sara Dizzell; Resources, Eileen Hutton, Katherine Morrison and Jennifer Stearns; Supervision, Eileen Hutton and Katherine Morrison; Visualization, Chiara-Maria Homann; Writing – original draft, Chiara-Maria Homann, Sara Dizzell and Jennifer Stearns; Writing – review & editing, Chiara-Maria Homann, Sandi Azab and Jennifer Stearns.

## Institutional Review Board Statement

The study was conducted according to the guidelines of the Declaration of Helsinki and was approved by the Research Ethics boards at Hamilton Integrated Research Ethics Board (12–201), St. Joseph’s Healthcare Hamilton (12–3721), Joseph Brant Hospital (000–022–14), Niagara Health System (2014–12–001) and Brant Community Healthcare System.

## Informed Consent Statement

Written informed consent was obtained from caregiver of the subject involved in the study.

## Data Availability Statement

The data presented in this study are available on request from the corresponding authors (J.C.S.). The data are not publicly available due to the potentially identifiable nature of the data and privacy concerns by study participants.

## Acknowledgments

The members of the GI-MDH Consortium are: John Penders^1,2,3^, Monique Mommers^4^, Alison C. Holloway^5^, Helen McDonald^6^, Elyanne M. Ratcliffe^7,8^, Jonathan D. Schertzer^8,9^, Mike G. Surette^8,9^, Lehana Thabane^10^, Susanne Lau^11^, Eckard Hamelmann^12^

^1^ School of Nutrition and Translational Research in Metabolism (NUTRIM), Department of Medical Microbiology, Maastricht University Medical Centre, Maastricht, the Netherlands.

^2^ in Vivo Planetary Health: an affiliate of the World Universities Network (WUN), West New York, New Jersey, USA.

^3^ School for Public Health and Primary Care (CAPHRI), Department of Medical Microbiology, Maastricht University Medical Centre, Maastricht, the Netherlands.

^4^ Department of Epidemiology, Care and Public Health Research Institute (CAPHRI), Maastricht University, Maas-tricht, The Netherlands.

^5^ Department of Obstetrics and Gynecology, McMaster University, Hamilton, ON, Canada.

^6^ McMaster Midwifery Research Centre, McMaster University, Hamilton, ON, Canada.

^7^ Department of Pediatrics, McMaster University, Hamilton, ON, Canada.

^8^ Farncombe Family Digestive Health Research Institute, McMaster University, Hamilton, Canada

^9^ Department of Biochemistry & Biomedical Sciences, McMaster University, Hamilton, Canada

^10^ Department of Clinical Epidemiology & Biostatistics, McMaster University, Hamilton, Canada.

^11^ Department of Pediatric Pulmonology, Immunology and Intensive Care Medicine, Charité Universitätsmedizin Berlin, Germany.

^12^ Children’s Center Bethel, Protestant Hospital Bethel, University of Bielefeld, Germany.

We would like to acknowledge the Baby & Mi research team for study coordination and data collection: Jenifer Li^1,2^, Julia Simioni^1,2^, Elizabeth Gunn^3^.

^1^ Department of Obstetrics and Gynecology, McMaster University, Hamilton, ON, Canada.

^2^ McMaster Midwifery Research Centre, McMaster University, Hamilton, ON, Canada.

^3^ Department of Pediatrics, McMaster University, Hamilton, ON, Canada.

We would also like to thank TMIC Alberta and TMIC Victoria for the processing of the metabolomic samples. Lastly, the authors would like to sincerely thank the participating caregiver, who committed to completing the requirements of this study, as well as McMaster Children’s Hospital and McMaster University.

