## Supplementary Material for "Investigation of the impact of stool collection methods on the metabolomics analysis/profiles of infant fecal samples"

### Solid Stool versus Diaper Liner: Stool Collection Method Impact Concentrations of Metabolites Recovered from Infant Stool Samples.

#### Supplementary Materials

**Table S1:** CV for metabolites measured in stool and liner with NMR.

| Metabolite | CV Solid Stool (%) | CV Stool Saturated Liner (%) |
|---|---|---|
| 2-Hydroxybutyrate | 2.25 | 3.98 |
| 2-Hydroxyisobutyrate | 6.28 | 9.55 |
| 4-Aminobutyrate | 1.65 | 3.64 |
| 4-Hydroxyphenylacetate | 1.28 | 3.60 |
| 5-Aminopentanoate | 2.86 | 0.49 |
| Acetate | 4.40 | 1.12 |
| Acetone | 3.38 | 4.98 |
| Alanine | 2.16 | 0.99 |
| Asparagine | 1.59 | 2.63 |
| Aspartate | 1.01 | 2.93 |
| Betaine | 4.41 | 5.48 |
| Butyrate | 4.08 | 3.74 |
| Cadaverine | 1.34 | 1.99 |
| Carnitine | 2.47 | 2.24 |
| Choline | 2.77 | 1.56 |
| Citrate | 2.59 | 2.12 |
| Creatine | 5.54 | 4.82 |
| Creatinine | 12.40 | 6.56 |
| Dimethylamine | 10.76 | 16.55 |
| Ethanol | 0.18 | 2.79 |
| Formate | 7.11 | 4.92 |
| Fucose | 0.67 | 0.85 |
| Fumarate | 1.40 | 5.27 |
| Galactose | 2.31 | 3.02 |
| Glucose | 1.54 | 0.98 |
| Glutamate | 1.44 | 0.68 |
| Glutamine | 1.26 | 1.64 |
| Glycerol | 0.06 | 0.20 |
| Glycine | 0.30 | 0.21 |
| Hypoxanthine | 1.46 | 3.02 |
| Isobutyrate | 3.11 | 0.93 |
| Isocaproate | 131.73 | 7.66 |
| Isoleucine | 1.20 | 0.77 |
| Isovalerate | 1.71 | 1.05 |
| Lactate | 2.11 | 3.54 |
| Leucine | 0.39 | 2.46 |
| Lysine | 0.81 | 2.57 |

| | | |
|---|---|---|
| Malate | 0.80 | 1.70 |
| Malonate | 0.18 | 0.42 |
| Methanol | 0.94 | 2.88 |
| Methylamine | 11.79 | 11.29 |
| N6-Acetyllysine | 2.22 | 2.13 |
| O-Phosphocholine | 3.52 | 4.88 |
| Proline | 2.95 | 4.23 |
| Propionate | 2.07 | 0.36 |
| Propylene glycol | 1.22 | 4.03 |
| Putrescine | 3.82 | 2.78 |
| Pyroglutamate | 1.97 | 5.26 |
| Pyruvate | 2.61 | 1.99 |
| Serine | 1.38 | 5.26 |
| Succinate | 2.17 | 5.81 |
| Taurine | 2.00 | 2.42 |
| Threonine | 2.42 | 5.80 |
| Trimethylamine | 6.07 | 7.15 |
| Tryptophan | 3.98 | 2.48 |
| Tyramine | 5.19 | 1.35 |
| Tyrosine | 0.64 | 0.54 |
| Uracil | 1.17 | 2.27 |
| Valerate | 2.72 | 1.58 |
| Valine | 2.41 | 1.35 |
| Xanthine | 3.33 | 2.62 |
| methionine | 3.59 | 1.08 |
| myo-Inositol | 3.85 | 1.10 |
| p-Cresol | 132.36 | 7.96 |
| phenylacetic acid | 2.39 | 1.91 |
| phenylalanine | 1.13 | 5.98 |
| β-Alanine | 4.71 | 4.70 |

**Table S2:** CV for metabolites measured in stool and liner with DI-MS.

| Metabolite | CV Solid Stool (%) | CV Stool Saturated Liner (%) |
|---|---|---|
| Creatinine | 3.75 | 5.88 |
| Glycine | 3.20 | 5.93 |
| Alanine | 10.80 | 9.15 |
| Serine | 6.28 | 9.82 |
| Proline | 6.63 | 10.15 |
| Valine | 6.56 | 9.39 |
| Threonine | 9.66 | 9.16 |
| Taurine | 13.22 | 0.54 |
| Putrescine | 8.20 | 2.60 |
| trans-hydroxy-Proline | 2.69 | 12.07 |
| Leucine | 3.83 | 6.62 |

| | | |
|---|---|---|
| Isoleucine | 7.96 | 6.48 |
| Asparagine | 9.08 | 8.26 |
| Aspartic acid | 20.77 | 5.44 |
| Glutamine | 7.21 | 5.12 |
| Glutamic acid | 6.11 | 17.48 |
| Methionine | 5.25 | 8.24 |
| Histidine | 4.43 | 9.65 |
| alpha-Aminoadipic acid | 93.82 | 8.88 |
| Phenylalanine | 8.96 | 7.48 |
| Methionine-sulfoxide | 8.32 | 6.81 |
| Arginine | 15.97 | 10.06 |
| Acetyl-Ornithine | 6.54 | 13.73 |
| Citrulline | 20.77 | 27.13 |
| Serotonin | 47.92 | 11.71 |
| Tyrosine | 4.18 | 12.66 |
| Asymmetric dimethylarginine | 14.33 | 4.01 |
| Total dimethylarginine | 10.82 | 15.61 |
| Tryptophan | 11.14 | 8.06 |
| Ornithine | 7.33 | 5.82 |
| Lysine | 12.61 | 16.75 |
| Spermidine | 1.28 | 12.04 |
| Spermine | 15.18 | 4.23 |
| Sarcosine | 14.03 | 21.70 |
| Diacetylspermine | 8.11 | 8.26 |
| Creatine | 14.66 | 8.81 |
| Betaine | 4.67 | 11.81 |
| Choline | 6.50 | 7.76 |
| Methylhistidine | 8.12 | 8.12 |
| LYSOC14:0 | 19.10 | 2.37 |
| LYSOC16:1 | 34.73 | 51.73 |
| LYSOC16:0 | 48.01 | 23.07 |
| LYSOC17:0 | 15.74 | 77.68 |
| LYSOC18:2 | 32.15 | 44.53 |
| LYSOC18:1 | 64.20 | 12.67 |
| LYSOC18:0 | 38.84 | 20.41 |
| LYSOC20:4 | 12.30 | 21.29 |
| LYSOC20:3 | 9.93 | 11.75 |
| LYSOC24:0 | 28.34 | 20.35 |
| LYSOC26:1 | 48.43 | 37.18 |
| LYSOC26:0 | 47.67 | 22.15 |
| LYSOC28:1 | 52.11 | 5.10 |
| LYSOC28:0 | 10.62 | 18.16 |
| 14:1SMOH | 18.44 | 32.23 |
| 16:1SM | 36.55 | 23.63 |
| 16:0SM | 51.32 | 27.59 |
| 16:1SMOH | 22.10 | 10.77 |

| | | |
|---|---|---|
| 18:1SM | 27.69 | 51.42 |
| PC32:2AA | 39.48 | 16.60 |
| 18:0SM | 21.19 | 7.69 |
| 20:2SM | 11.98 | 36.20 |
| PC36:0AE | 52.50 | 1.99 |
| PC36:6AA | 28.57 | 5.45 |
| PC36:0AA | 22.25 | 15.22 |
| 22:2SMOH | 12.05 | 22.77 |
| 22:1SMOH | 48.97 | 23.32 |
| PC38:6AA | 67.05 | 47.04 |
| PC38:0AA | 13.11 | 14.58 |
| PC40:6AE | 15.81 | 23.52 |
| 24:1SMOH | 27.62 | 38.51 |
| PC40:6AA | 31.28 | 40.74 |
| PC40:2AA | 20.54 | 25.79 |
| PC40:1AA | 5.67 | 15.21 |
| C0 | 9.61 | 3.65 |
| C2 | 14.01 | 8.87 |
| C3:1 | 16.27 | 8.19 |
| C3 | 16.88 | 11.79 |
| C4:1 | 8.58 | 1.20 |
| C4 | 21.41 | 7.07 |
| C3OH | 20.14 | 13.24 |
| C5:1 | 13.24 | 3.37 |
| C5 | 15.63 | 13.75 |
| C4OH | 9.92 | 2.36 |
| C6:1 | 5.96 | 2.76 |
| C6 | 19.00 | 4.78 |
| C5OH | 10.84 | 8.57 |
| C5:1DC | 15.56 | 11.86 |
| C5DC | 40.20 | 11.43 |
| C8 | 21.27 | 10.79 |
| C5MDC | 16.88 | 14.49 |
| C9 | 6.35 | 6.53 |
| C7DC | 12.74 | 7.17 |
| C10:2 | 11.71 | 4.79 |
| C10:1 | 13.57 | 4.19 |
| C10 | 12.29 | 4.22 |
| C12:1 | 13.70 | 1.89 |
| C12 | 13.18 | 6.26 |
| C14:2 | 2.10 | 16.24 |
| C14:1 | 16.84 | 10.57 |
| C14 | 14.99 | 6.23 |
| C12DC | 10.38 | 1.52 |
| C14:2OH | 3.81 | 5.63 |
| C14:1OH | 14.62 | 15.92 |

| | | |
|---|---|---|
| C16:2 | 9.70 | 3.92 |
| C16:1 | 15.68 | 10.44 |
| C16 | 20.46 | 9.59 |
| C16:2OH | 8.10 | 16.68 |
| C16:1OH | 13.64 | 6.11 |
| C16OH | 9.00 | 4.86 |
| C18:2 | 8.78 | 3.98 |
| C18:1 | 15.47 | 2.16 |
| C18 | 5.88 | 13.56 |
| C18:1OH | 20.72 | 7.10 |
| Glucose | 7.41 | 19.20 |

**Table S3:** CV for metabolites measured in stool and liner with UPLC-MS.

| Metabolite | CV Solid Stool (%) | CV Stool Saturated Liner (%) |
|---|---|---|
| Cholic acid | 14.73 | 20.44 |
| Deoxycholic acid | 7.10 | 15.01 |
| Lithocholic acid | 14.42 | 15.53 |
| Allocholic acid | 15.05 | 17.18 |
| Chenodeoxycholic acid | 12.89 | 22.26 |
| Dehydrocholic acid | 11.11 | 9.09 |
| Dehydrolithocholic acid | 6.19 | 17.32 |
| 7-Ketodeoxycholic acid | 14.92 | 17.54 |
| 7-Ketolithocholic acid | 9.96 | 20.10 |
| Apocholic acid | 7.41 | 17.17 |
| Hyodeoxycholic acid | 44.30 | 40.00 |
| Murocholic acid | 14.43 | 25.00 |
| Ursodeoxycholic acid | 12.35 | 22.54 |
| Dioxolithocholic acid | 18.03 | 15.26 |
| 12-Ketochenodeoxycholic acid | 21.75 | 12.07 |
| 3-Oxocholic acid | 14.23 | 15.03 |
| α-Muricholic acid | 17.47 | 28.07 |
| β-Muricholic acid | 21.32 | 21.39 |
| λ-muricholic acid | 14.00 | 22.82 |
| Ш-muricholic acid | 13.59 | 15.74 |
| Ursocholic acid | 14.55 | 15.11 |
| Norcholic acid | 15.56 | 20.14 |
| Glycochenodeoxycholic acid | 24.29 | N/A |
| Glycocholic acid | 6.54 | 9.36 |
| Taurochenodexycholic acid | 31.11 | 14.70 |
| Taurocholic acid | 10.78 | 11.85 |
| Taurohyocholic acid | 14.32 | 8.66 |
| Tauro-α-muricholic acid | 10.18 | 5.88 |
| Tauro-Ш-muricholic acid | 10.26 | 23.15 |

| | | |
|---|---|---|
| Deoxycholic acid-3-glucuronide | 37.55 | 11.55 |
| Lithocholic acid-3-sulfate | 8.18 | 10.97 |
| Deoxycholic acid-3-sulfate | 23.28 | 17.85 |
| Allocholic acid-3-sulfate | 16.93 | 16.58 |
| Cholic acid-3-sulfate | 18.92 | 16.55 |
| Glycochenodeoxycholic acid-3-sulfate | 15.57 | 15.56 |
| Taurochenodeoxycholic acid-3-sulfate | 17.28 | 12.96 |
| Tauroursodeoxycholic acid-3-sulfate | 26.36 | 32.73 |
| Chenodeoxycholic acid-3-sulfate | 13.45 | 18.99 |
| Ursodeoxycholic acid-3-sulfate | 7.73 | 18.94 |

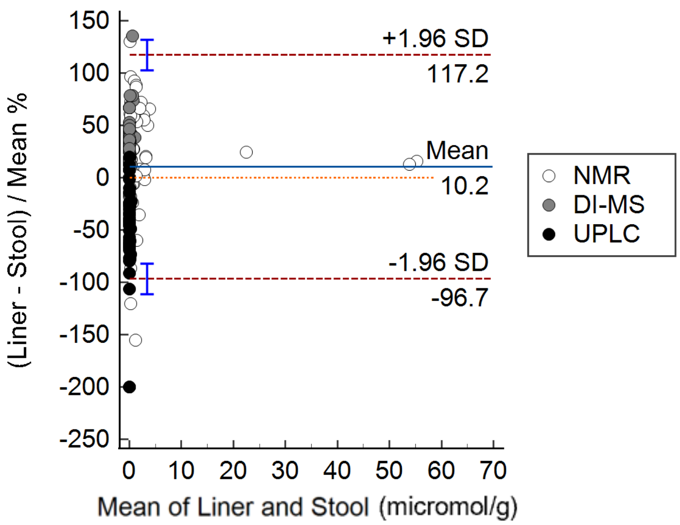


**Figure S1:** Bland-Altman plot comparing metabolite concentrations between methods for stool collection (solid stool vs. liner), colored by metabolic platform. Metabolites outside of the confidence intervals of the upper and lower limits are considered different between methods.

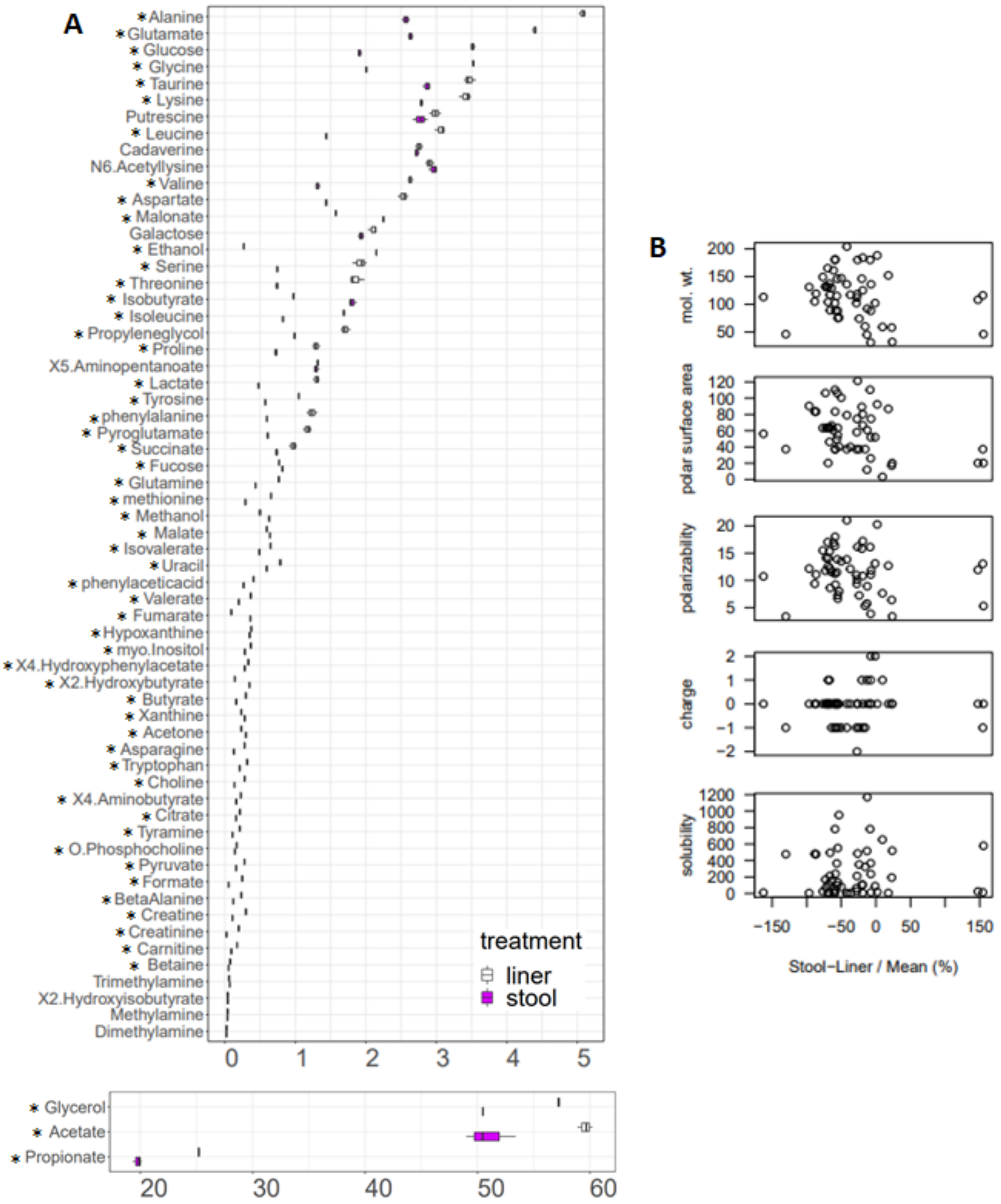


**Figure S2:** Metabolite, including short chain fatty acid, concentrations (μmol/g) measured in stool and liner with NMR. (**A)** Boxplot of metabolite concentrations, * = significant differences between stool sample collection methods, after adjustment for multiple testing. (**B)** Characteristics of the metabolites in relation to their log2 fold change by sample collection method.

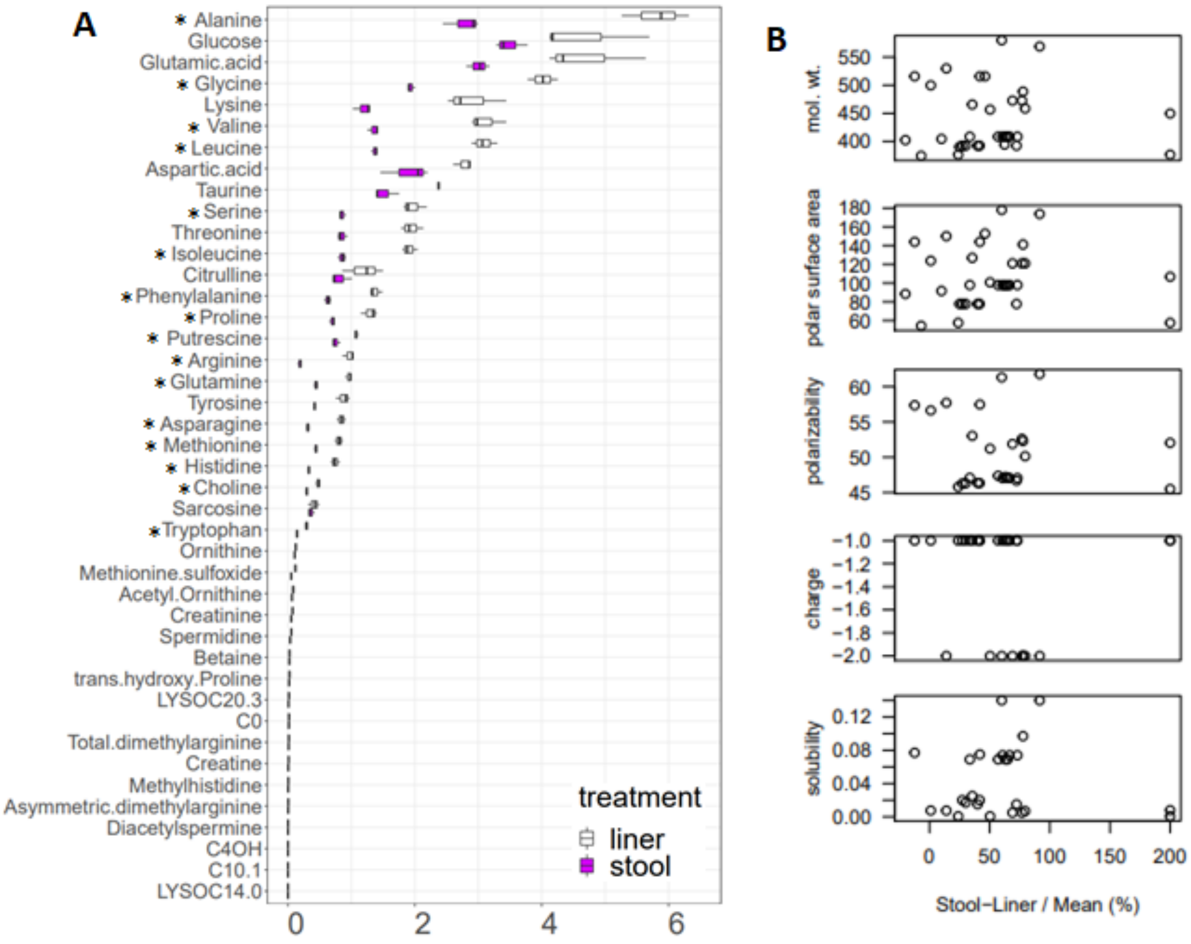


**Figure S3.** Metabolite concentrations (μmol/g) measured in stool and from liner with DI-MS/MS. (**A)** Boxplot of metabolite concentrations, * = significant differences between stool sample collection methods, after adjustment for multiple testing. (**B)** Characteristics of the metabolites in relation to their log2 fold change by sample collection method.

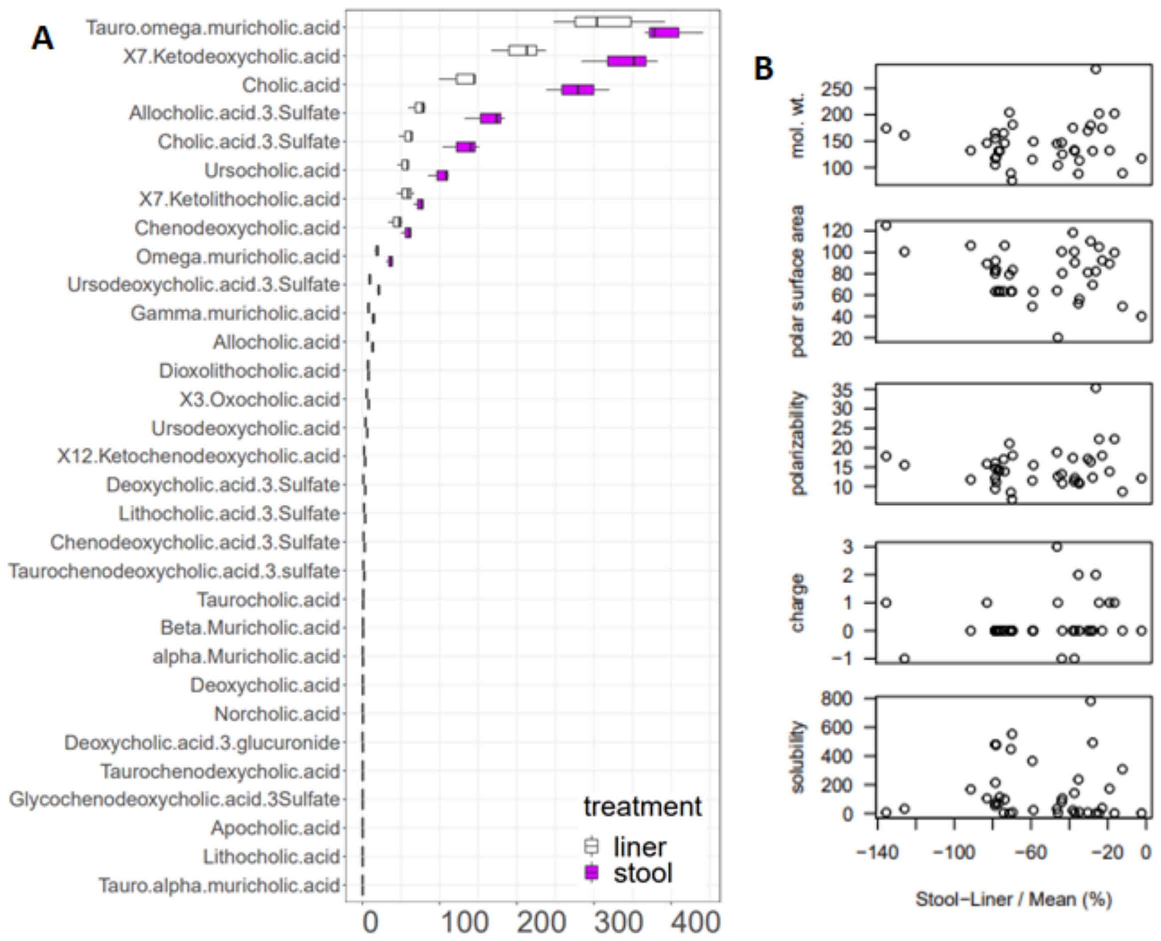


**Figure S4.** Bile acid concentration (nmol/g) measured in stool and from liner with UPLC-MS. (**A)** Boxplot of metabolite concentrations. (**B)** Characteristics of the metabolites in relation to their log2 fold change by sample collection method.
